# The Evaluative Role of Rostrolateral Prefrontal Cortex in Rule-Based Category Learning

**DOI:** 10.1101/107110

**Authors:** Dmitrii Paniukov, Tyler Davis

**Affiliations:** Department of Psychological Sciences, Texas Tech University

**Keywords:** category learning, rostrolateral prefrontal cortex, rule evaluation, rule switching, fMRI

## Abstract

Category learning is a critical neurobiological function that allows organisms to simplify a complex world. Rostrolateral prefrontal cortex (rlPFC) is often active in neurobiological studies of category learning; however, the specific role this region serves in category learning remains uncertain. Previous category learning studies have hypothesized that the rlPFC is involved in switching between rules, whereas others have emphasized rule abstraction and evaluation. We aimed to clarify the role of rlPFC in category learning and dissociate switching and evaluation accounts using two common types of category learning tasks: matching and classification. The matching task involved matching a reference stimulus to one of four target stimuli. In the classification task, participants were shown a single stimulus and learned to classify it into one of two categories. Matching and classification are similar but place different demands on switching and evaluation. In matching, a rule can be known with certainty after a single correct answer. In classification, participants may need to evaluate evidence for a rule even after an initial correct response. This critical difference allows isolation of evaluative functions from switching functions. If the rlPFC is primarily involved in switching between representations, it should cease to be active once participants settle on a given rule in both tasks. If the rlPFC is involved in rule evaluation, its activation should persist in the classification task, but not matching. The results revealed that rlPFC activation persisted into correct trials in classification, but not matching, suggesting that it continues to be involved in the evaluations of evidence for a rule even after participants have arrived at the correct rule.

**Highlights:**

- Differences between rule-based matching and classification tasks were highlighted.
- Rostrolateral prefrontal cortex is involved in evaluation of evidence for a rule in rule-based category learning tasks.

## 1. Introduction

Friend or foe? Predatory or prey? Edible or poisonous? Category learning is a fundamental cognitive capacity that is critical for survival. Grouping objects into categories allows organisms to generalize information to novel examples and make inferences about their characteristics. For example, having the category *bird* can help a person identify new species they have never seen before as birds, and make predictions about their biological features. As a complex cognitive function, many brain regions are involved in category learning including the prefrontal cortex (PFC), medial temporal lobes, striatum, and visual cortex (Ashby & Maddox, 2005, 2011; Poldrack & Foerde, 2008; Seger & Miller, 2010; Smith & Grossman, 2008).

Although there are many types of category learning, one of the most studied types in cognitive neuroscience is rule-based category learning. In rule-based category learning, people learn a logical rule that can be used to determine whether items are members of the category or not. Many real world categories are associated with logical, albeit imperfect, rules. For example, members of the category *bird* can often be categorized based on whether they fly and lay eggs. Broadly speaking, rule-based categorization is thought to depend upon executive cortico-striatal loops that connect the PFC and the head of the caudate nucleus (Alexander et al., 1986; Seger & Miller, 2010). Although early work with patients and fMRI tended to treat the PFC as being involved in executive functions as a whole (Konishi et al., 1998, Robinson et al., 1980), recent work in cognitive neuroscience has begun to test whether specific subregions of the PFC serve distinct mechanisms (Seger, 2008; Seger & Miller, 2010; Ma et al., 2016).

One critical PFC region that has thus far eluded a thorough explanation in terms of its role in category learning is the lateral parts of the fronto-polar cortex, also known as the rostrolateral prefrontal cortex (rlPFC). The rlPFC is known to be involved in a broad array of higher-level cognitive functions including abstract symbolic reasoning and analogical problem solving (Green et al., 2006; Specht et al., 2009), relational category learning (Davis et al., 2017), and goal-directed reward learning (Spreng et al., 2010). It is often described as a seat of human reasoning powers as it is significantly larger in humans than other primates, and its development across childhood tracks the development of fluid reasoning capacities (Gogtay et al., 2004; Semendeferi et al., 2011).

In humans, increased rlPFC activation is often observed for rule-based tasks involving abstract symbolic (Specht et al., 2009) or relational reasoning (Davis et al., 2017; Gray et al., 2003; Wendelken & Bunge, 2009). The rlPFC’s precise mechanistic role in these tasks has been described in a number of different ways. One characterization of the rlPFC focuses on switching between representations. For example, the rlPFC has been found to be more active when participants switch between cognitive sets in rule-based tasks (Konishi et al., 1998, 2002; Monchi et al., 2001; Strange et al., 2001; Liu et al., 2015), in tasks requiring switching between internally focused and externally focused attention (Burgess et al., 2007), and in reward learning tasks when exploring the values of different choices (Daw et al., 2006).

Other studies of rlPFC function have focused on its role in forming abstract symbolic rule representations and testing and evaluating such rules (Badre & D'Esposito, 2009; Vendetti & Bunge, 2014; Wendelken et al., 2012). Classic examples of the rlPFC’s involvement in symbolic rule use include its stronger engagement for rules requiring higher-order relational integration (rules that integrate over multiple lower-order relationships) in tasks akin to Raven’s progressive matrices (Christoff et al., 2001; Kroger et al., 2002), and tasks requiring participants to answer whether such a higher order relation is present in a stimulus (Bunge et al., 2009; Nee et al., 2014). More recently, the rlPFC has also been found to be engaged in the incremental learning of rules across trials (Badre et al., 2010; Davis et al., 2017). In these cases, the rlPFC may not only be integrating relationships within stimuli to form rules, but also integrating or accumulating information across trials to evaluate evidence for a rule and test how it applies to new stimuli.

Indeed, in Davis and colleagues (2017), the rlPFC was active early in learning as participants acquired a new relational rule, but then later only on trials in which they needed to apply the rule to novel stimuli. This evaluative role in integrating and accumulating information for a rule potentially connects the rlPFCs role in rule learning with recent findings of its engagement during meta-cognitive judgments (Fleming et al., 2012), and may explain why the rlPFC is often active during learning of rules that do not strictly involve higher-order relational integration within a stimulus or trial (e.g., Seger & Cincotta, 2006; Liu et al., 2015).

To test the distinction between rule switching and evaluation accounts of rlPFC function, we compared activation during two commonly used rule-based tasks from the category learning literature — matching and classification tasks. Matching tasks, like the Wisconsin Card Sorting Test (Heaton, 1993), involve matching a multidimensional reference stimulus to a set of target stimuli that each match the reference on a single dimension. There is a rule that determines which target to match the reference to that is based on the stimulus dimensions. Participants learn the matching rule through trial and error. For example, in the matching task that we used in the present study, reference stimuli were schematic beetles that differed in terms of their legs, tail, antennae, and mandibles. Participants learned to match these reference stimuli to target beetles by choosing different candidate targets and receiving feedback. Often matching tasks will cycle through a number of rules forcing participants to abandon rules and shift to a new rule when old rules cease to be useful.

As a neuropsychological measure, the primary process of interest in matching tasks like the Wisconsin Card Sorting Test is the process of shifting between cognitive sets to accommodate novel rules and suppress the previously correct rules. Consistent with the theory that the rlPFC governs representational switching, results from a number of neuroimaging studies have identified activation in rlPFC during trials in which the rule is switched (Konishi et al., 1998, 2002; Monchi et al., 2001; Strange et al., 2001; Liu et al., 2015). However, switch trials as well as the immediately following trials in which the novel rule is being acquired also place demands upon rule evaluation mechanisms. Participants must not only switch between the previous and new candidate rule representations, but also begin to test and evaluate evidence for new candidate rules. One critical aspect of simple matching tasks, like the Wisconsin Card Sorting Test, is that participants gain some conclusive information to evaluate a rule on every trial: if a choice is wrong, a candidate rule can be eliminated; if a choice is correct, the rule is known with certainty. This aspect of matching tasks means that evaluation of evidence for a rule and switching are strongly intertwined because participants will switch rules whenever they are wrong (Konishi et al., 1998; Monchi et al., 2001) and quit evaluating new information once they are correct. Thus, for participants performing rationally, every possible switch trial is also a trial in which new information is being obtained and evaluated. Likewise, every trial that provides new information about the rule is a trial in which the participant has switched the rule they are using.

We contrast matching tasks with another common type of rule-based category learning: classification tasks. In classification tasks, participants learn a rule that allows them to sort multidimensional stimuli into two or more categories based on trial and error. When the rule is based on a single dimension, classification tasks involve very similar mechanisms to single-dimension matching tasks like the Wisconsin Card Sorting Test. Participants test candidate rules, update their candidate rules based on feedback, and maintain rules in working memory once they have arrived at them. Indeed, because of this overlap in processing requirements, many neurobiological theories of category learning assume that matching and classification tasks have strong overlap in terms of the systems involved in acquiring and storing new rules and use results from both types of tasks interchangeably (e.g., Ashby & Maddox, 2005). However, classification tasks differ from matching tasks critically in requiring more extensive cross-trial evaluation of evidence for a rule, while otherwise keeping the demands on representational switching constant.

In classification tasks, participants are shown single examples of stimuli and learn, using trial-and-error, to classify the stimuli into one or more categories based on the stimulus dimensions. Psychologically, as in matching tasks, participants must evaluate evidence for the correct rule by switching their attention between different stimulus dimensions to test various candidate rules across trials. Also, like matching tasks, it is possible to eliminate a particular rule based on feedback in a single trial — if a participant tries the rule “thick legs = Category A”, incorrect feedback will tell them that they need to switch the rule they are using and eliminate this rule. However, individual correct trials in classification tasks contain less information about the correct rule than in matching tasks. From an optimal observer standpoint, a single trial in a matching task can tell whether a rule is correct or incorrect, however, even when behaving optimally, on average, even several correct trials in a classification task can leave potential candidate rules to decide amongst (see Figure 1 for an illustration of evaluative demands in the tasks). This optimal observer perspective suggests that some intrinsic uncertainty about the rule should remain even after participants begin getting trials correct.

**Figure 1.**
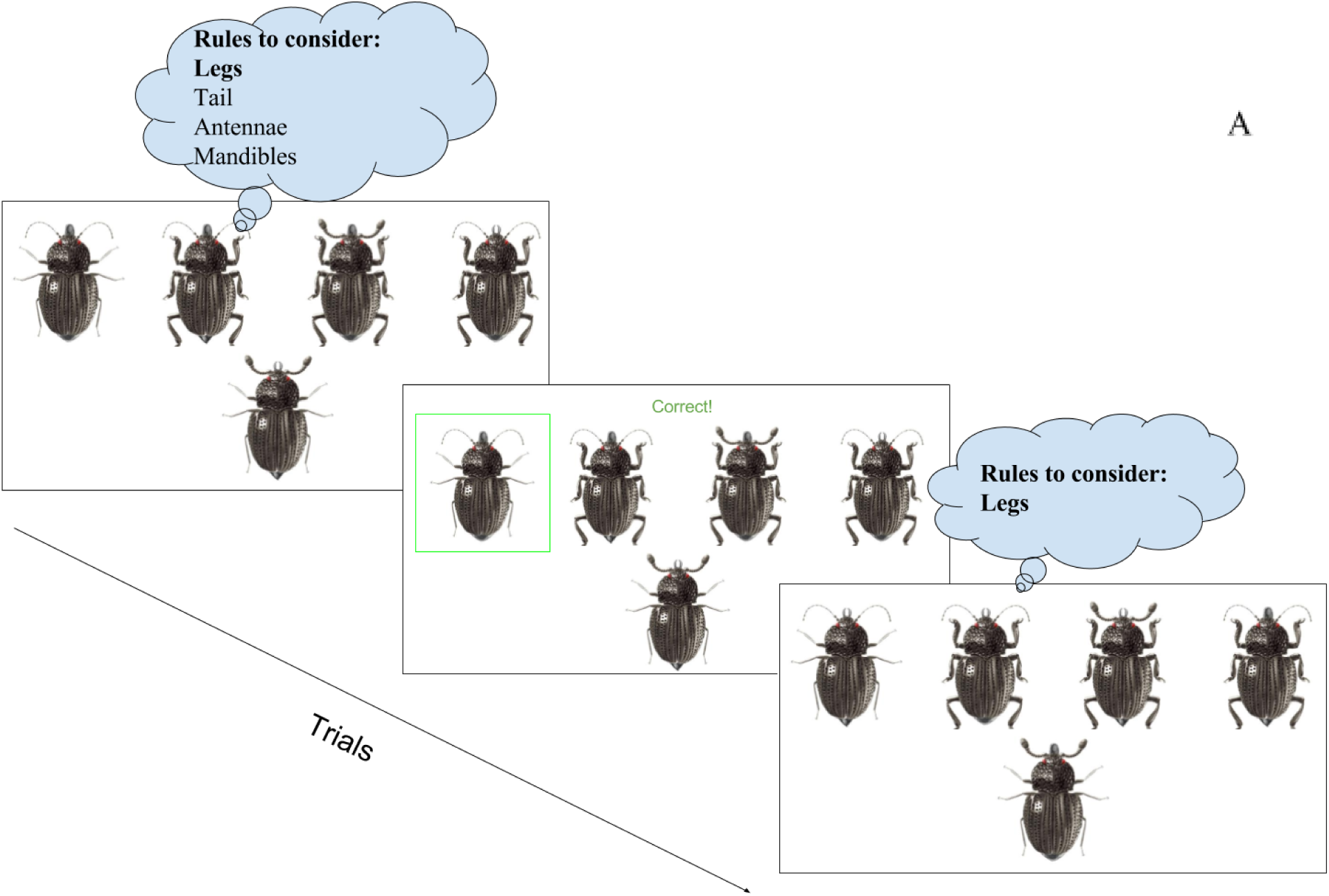
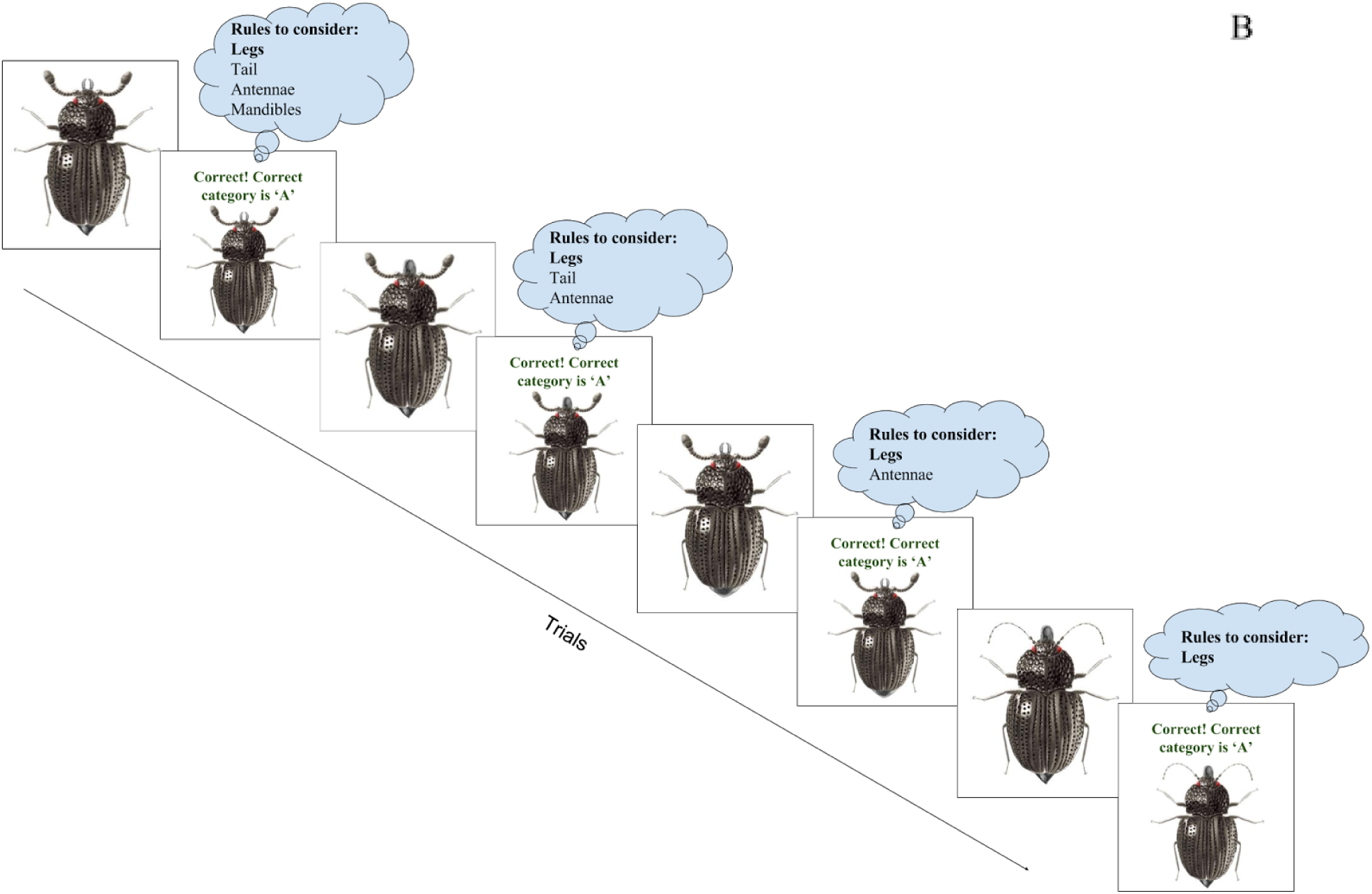
Examples of matching (A) and classification tasks (B) and how rules are eliminated in both tasks. In matching tasks, when the rule is correctly selected, all other rules are eliminated. Contrastingly, in classification tasks, even if a correct rule is selected initially, from an optimal observer’s standpoint, several additional trials may be necessary to rule out other rules that are possible given the history of stimulus-category pairings participants have seen up to that point. In both of these cases, participants have arrived at the correct rule dimension (“legs” in bold) on the first trial of the depicted sequence. In matching, a single instance of correct feedback establishes that “legs” are the rule-dimension. In classification, although the participant may start out using the legs rule, they may only fully eliminate other possible rules that are consistent with the stimulus history after a number of correct trials. For example, if in the first trial the participant chooses category ‘A’ for the thin legged beetle and gets correct feedback, all rules will still be under consideration. When an optimal observer chooses the same category for the thin legged beetle on the second trial and receives correct feedback, only the rules “pointy tail = ’A’; thick antennae = ‘A’; and bisected mandibles = ‘A’” become active. They only become eliminated after the optimal observer encounters variations of the “thin legs = ‘A’” rule, in which these candidate rules do not also hold.

Of course, participants in classification tasks rarely perform like an optimal observer. Instead, extensive research suggests that participants in classification tasks use a win-stay-lose-shift strategy where they will not switch from a rule when they are getting correct feedback and will only switch to remaining candidate rules when they get negative feedback (Shepard et al., 1961; Nosofsky, Palmeri, and McKinley, 1993; Wilson & Niv, 2011; Niv et al., 2015). Likewise, they will not tend to eliminate rules based on all of the information they have encountered (as the optimal observer in Figure 1 does), and instead will eliminate rules sequentially. Together with the optimal observer perspective, participants’ tendencies to not switch rules while they are getting trials correct suggests a critical asymmetry between matching tasks and classification tasks that can be harnessed to isolate evaluation processes from switching. Specifically, because uncertainty will remain regarding the true rule even after participants begin getting trials correct, but subjects will not switch rules while getting items correct, we can isolate trials in which they continue to evaluate evidence for a rule but do not switch between rules.

To test how differences in evaluation demands between matching and classification tasks impact rlPFC activation, we had participants complete the matching and classification tasks using schematic beetle stimuli. In both tasks, participants would iteratively learn rules via trial and error. Once rules were learned (defined by four correct trials in a row) the rule would switch and the participant would begin learning a new rule. For each rule there was therefore a *rule learning* phase, when participants were narrowing down and switching between rules, and a brief *rule application* phase, in which participants were using the final rule that they have arrived at (Figure 2).

**Figure 2.**
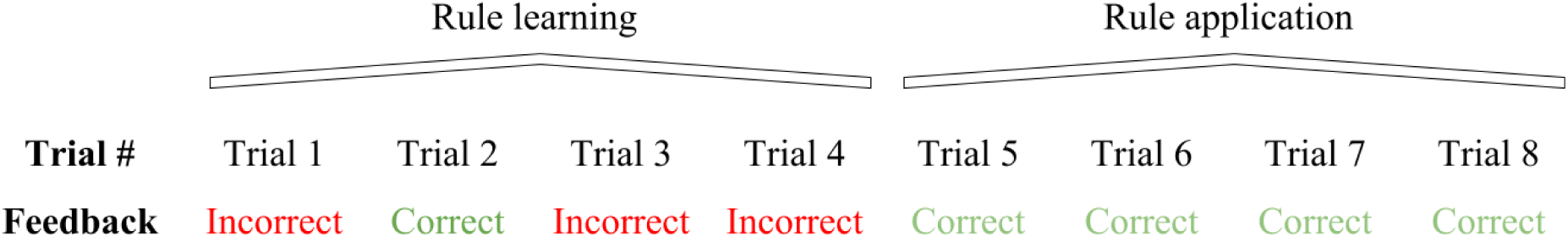
An example of how the rule learning and rule application phases were separated for analysis. The last four or more contiguous correct trials were labeled as rule application. In this phase participants applied a final rule they had learned. All trials before the final four contiguous correct trials were labeled as rule learning.

The different accounts of rlPFC function can be used to make predictions for the rule learning and application phases. If the rlPFC is primarily involved in switching, it should only be active during the rule learning phases of both matching and classification tasks, when participants are trying different rules and switching between them. Contrastingly, if the rlPFC is involved in evaluation, it should be active not only during the rule learning phase of both tasks, but also during the rule application phase for classification when participants have arrived at the correct rule, but are still evaluating evidence in support of it. Because rule evaluation is not necessary during matching once the correct rule is found, rlPFC should not be active during the application phase of the matching task.

To foreshadow our results, we found evidence consistent with the rule evaluation hypothesis. The rlPFC was more active in the classification task than the matching task during both the rule learning and rule acquisition phases, suggesting that the rlPFC’s role extends beyond when participants are switching between rules in rule-based category learning tasks.

## 2. Method

### 2.1. Participants

Twenty-seven participants were recruited from the Texas Tech University community via an electronically posted announcement. Participants were required to be at least 18 years of age, right-handed, have a minimum of an eighth grade education, speak English fluently, and not have any contraindications for MRI research. Participants were compensated with $35 for the study. Two participants fell asleep during their scanning session and therefore were removed from the further analyses. One participant opted out from the last two scanning runs, but the rest of their data were used in the analyses. The study was approved by Human Research Protection Program at Texas Tech University.

### 2.2. Stimuli and Procedure

Participants were asked to sign a consent form, MRI-safety checklist and complete a computer-based tutorial. The participants were informed that the rules would be based off of the four different features of the stimuli, about switching rules after several consequent correct trials (to minimize anticipation of a new rule, we did not tell them after how many trials the rule will be switched), and were told each rule would be based on a single feature. Participants completed example trials and a brief test to examine their understanding of instructions, and they were allowed to ask questions if they were confused about any procedures. Upon completion of the screening forms and the tutorial, participants were placed into the scanner. In the scanner, participants completed four runs of the matching task and four runs of the classification task in an order that was balanced across participants.

Two tasks, matching and classification, utilized sixteen images of schematic beetles representing all possible combinations of the following four binary feature dimensions: legs (thick or thin), mandibles (closed or bisected), antennae (fuzzy or dotted), and tail (pointy or round; see Figure 3 for two examples with opposite features). For both tasks, the sixteen stimuli were presented in sequential randomized blocks. In a randomized block, a participant would go through all sixteen stimuli in a random order before seeing any of the same stimuli again. With this randomization, it was possible for as many as all of the features to change from trial-to-trial, or as few as zero, if the last stimulus of a randomized block and the first stimulus of the next block were the same.

**Figure 3.**
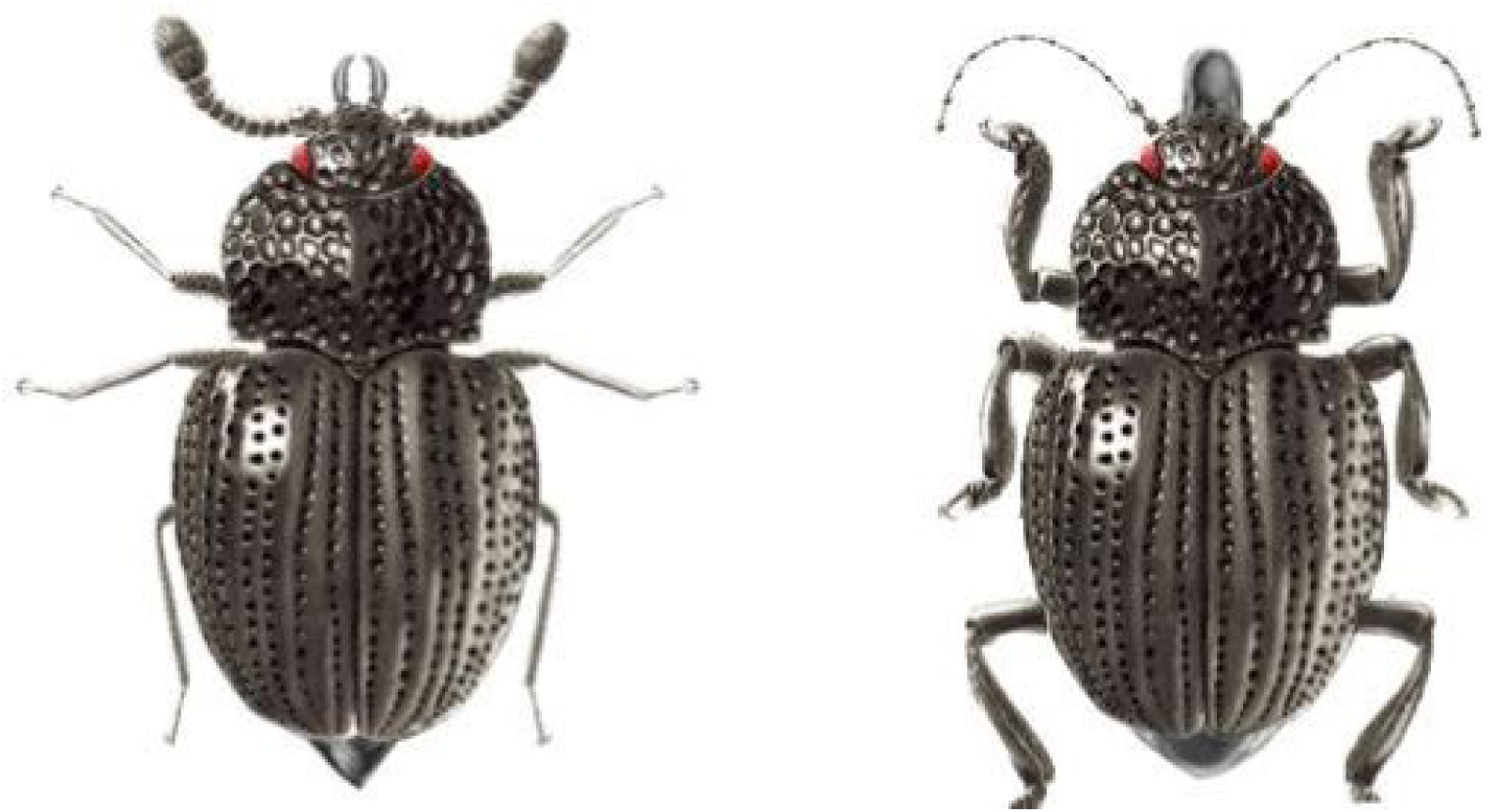
Stimuli of schematic beetles with completely distinct feature dimensions. The feature dimensions were legs (thick or thin), mandibles (closed or bisected), antennae (fuzzy or dotted), and tail (pointy or round).

In the matching task, each screen contained a reference beetle and four target beetles, each of which matched the reference beetle on a single dimension (see Figure 4). The position of the beetles on the screen was randomized to minimize effects of feature salience and balance motor responses.

**Figure 4.**
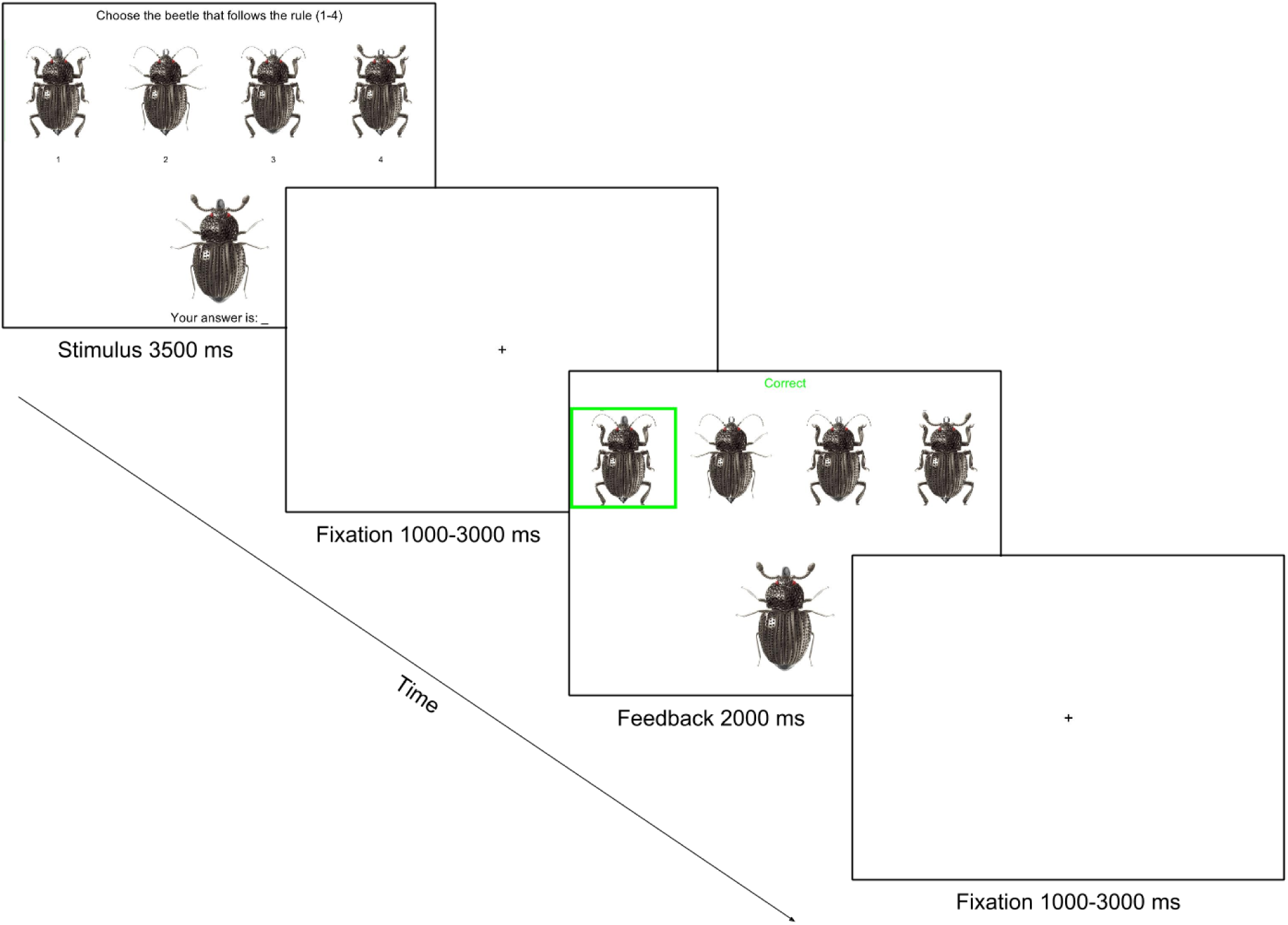
Example of the matching task. Participants saw a reference beetle on the bottom and four target beetles on top. The target beetles each matched the reference beetle on a single feature. Participants would select a target beetle and then receive feedback about whether their choice was correct or incorrect.

In the classification task, each screen contained a single beetle (see Figure 5). The beetles were assigned into category A or B, based on a random rule, defined by a single feature.

**Figure 5.**
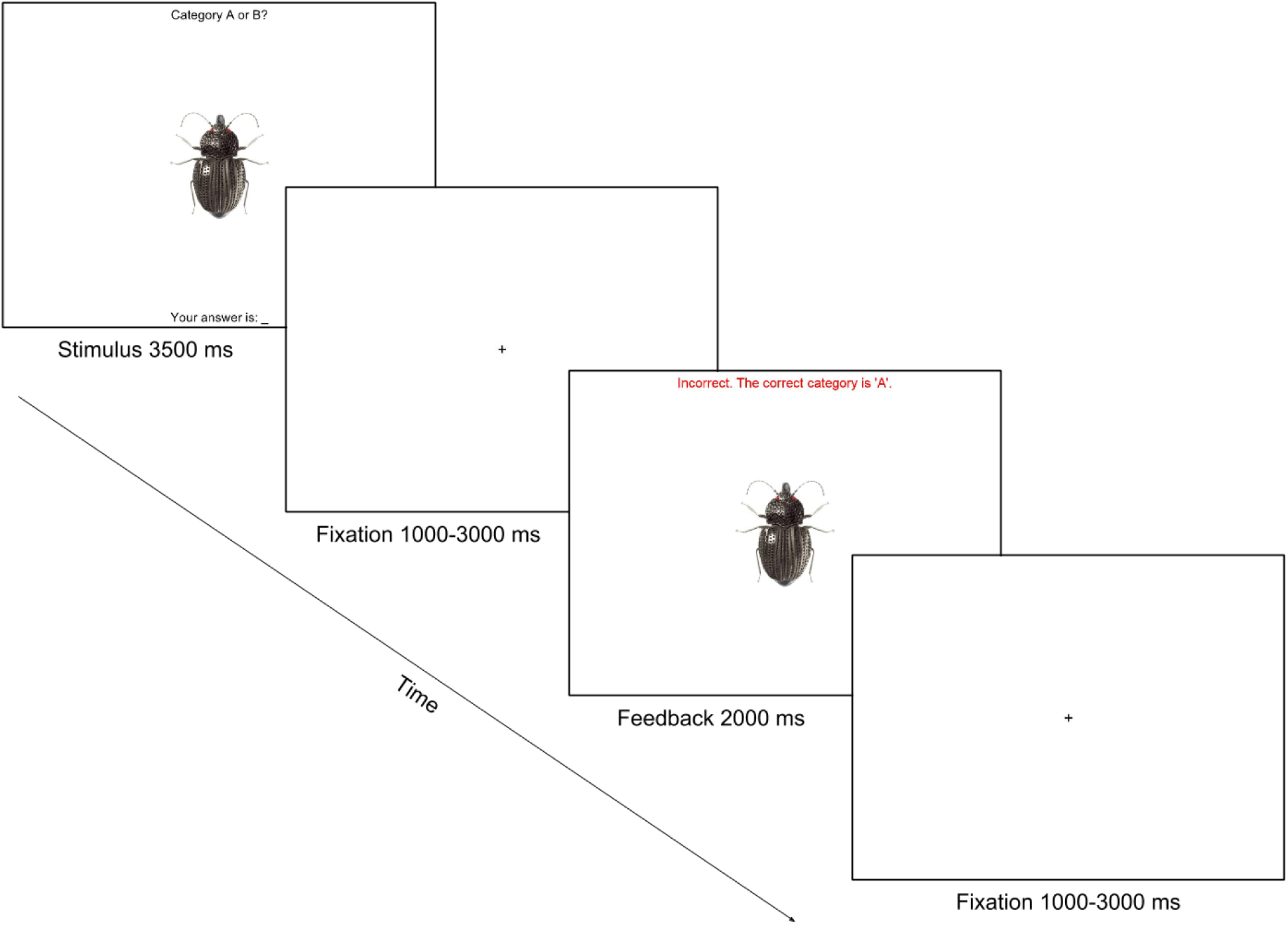
Example of the classification task. Participants saw a single beetle on the screen and were asked to categorize it as a member of Category A or B. After participants made the choice, they were given feedback about whether their choice was correct and the correct category label.

In both tasks, participants had 3.5 seconds to categorize a stimulus using a button box held in their right hand. After a brief fixation (1, 2, or 3s; mean = 2s), participants were provided with feedback for 2s, during which the beetle was presented again along with a message “Correct/Incorrect/Failed to respond” in the matching task or “Correct/Incorrect/Failed to respond.” and “The correct category is A/B” in the classification task. After the feedback, a brief fixation was presented (1, 2, or 3s; mean = 2s). Upon achieving a termination criterion of four correct answers in the row, the rule was switched to a new randomly selected rule. When a run was completed, participants were presented with a number of correctly solved rules during the run (i.e., “You successfully learned [number] categories! Good job!”). Participants had thirty-two trials in each run and eight runs total (four runs in a block for each task), followed by an anatomical scan. After the scan, participants were thanked, compensated and dismissed from the study.

### 2.3. fMRI Data Acquisition

The data were collected at Texas Tech Neuroimaging Institute using a Siemens Skyra 3T scanner with a 20-channel head coil. A T1-weighted sagittal MPRAGE was obtained with TR = 1.9s, TE = 2.49, flip angle = 9, matrix of 256x256, field of view = 240, slice thickness= 0.9 mm with a gap of 50%, 1 slice. T2-weighted BOLD echo planar images (EPI) were obtained with TR=2s, TE=30ms, flip angle = 90, matrix = 64x64, field of view = 192, slice thickness = 4 mm, 35 ascending axial slices, 156 volumes in each scanning run. The slice prescription was tilted off of parallel with AC/PC to reduce susceptibility artifact in orbital frontal cortex (Deichmann et al., 2003).

### 2.4. fMRI Data Preprocessing and Analysis

MRI data preprocessing included the following steps: DICOM imaging files were converted to NifTI files using dcm2nii from the Mricron software package (Rorden & Brett, 2000). Skulls were removed using the BET tool (Smith, 2002) from FSL software package (Jenkinson et al., 2012; Woolrich et al., 2009) for the BOLD EPI images and ANTs (Avants et al., 2009) with OASIS template (Avants & Tustison, 2014) for the T1 anatomical images. Motion correction was carried out using a 6-DOF affine transformation implemented in FSL’s MCFLIRT tool (Jenkinson et al., 2002). Data were smoothed with an 8 mm FWHM Gaussian kernel based on the standard criterion of two times the voxel size (Poldrack et al., 2011) and high-pass filtered (100s cut off). Finally, data were manually checked for quality issues such as visual spikes produced by the scanner, incorrect brain extraction, and excessive motion. The quality check revealed visual artifacts in the first run for four participants, and these runs were thus excluded from further analysis.

Functional MRI data analysis was carried out using a standard three-level analysis in FSL’s FEAT, as implemented in Nipype (Gorgolewski et al., 2011). The first-level analysis consisted of prewhitening using FSL’s FILM (Woolrich et al., 2001) to account for temporal autocorrelation, task-based regressors (see below) convolved with double-gamma hemodynamic response, and their temporal derivatives. Additional confound regressors of no interest included six motion parameters, their temporal derivatives and regressors for scrubbing high motion volumes exceeding a framewise displacement of 0.9mm (Siegel, et al., 2014). The same high pass filter setting that was used to process the fMRI data was used to process the design matrix. First-level statistical maps were registered to a standard space in a two-stage registration consisting of (1) registration of each BOLD timeseries to respective participants’ T1 MPRAGE using the BBR algorithm (Greve & Fischl, 2009) and (2) registration of the T1 to the standard space (MNI-152 brain template) using nonlinear ANTs registration (Avants et al., 2009). Second-level analysis combined across runs within a participant, and was carried out using a fixed effects model in FLAME (Beckmann et al., 2003; Woolrich et al., 2004; Woolrich, 2008). Second-level regressors included task type (matching vs. classification).

Third-level mixed effects analyses, treating participant as a random effect, examined whether first and second-level contrasts were significant across participants using a permutation test, implemented in FSL’s Randomise function (Winkler et al., 2014). This permutation test is a non-parametric approach to group-level analysis and multiple comparison correction that estimates the null distribution of cluster sizes from permutations of the data, alleviating recently publicized concerns about the accuracy of the parametric approximations underlying Gaussian Random Field Theory (Eklund et al. 2016). The final thresholding of statistical maps at *p*< 0.05 was done via a cluster mass correction (Bullmore et al., 2000), with a primary/cluster-forming threshold of *t*(24)=2.49, p < 0.01, one-tailed, and variance smoothing set at 8 mm. For a priori ROI analysis within the fronto-polar mask, we used the same cluster-mass based thresholding, but confined the analysis to the smaller volume.

All trials in both tasks were divided into either a rule learning phase in which participants were acquiring the correct rule or a rule application phase in which participants had acquired the rule and were applying it (see Figure 2 for an example of the phases for the classification task). In the matching task, all correct trials were considered as rule application trials due to the nature of the task, and all incorrect trials as rule learning. In the classification task only the last four or more continuous correct trials were considered as rule application, and the rest of the trials as rule learning. More than four correct trials in a row in which subjects used the same rule could occur in the classification task, even with the rule being programmed to switch after four correct. Specifically, due to our use of random blocks of stimuli, there were cases where the old rule could continue to work temporarily after a switch because stimuli would have features consistent with the same category response (A or B) in both the new and old rule. For example, if the previous rule was ‘thick legs = category A’ and the new rule was ‘thick antennae = category A’ then sometimes participants might see stimuli that had both of those dimensions, and thus the old rule would continue to work. Because participants would not know the rule had switched until they received negative feedback, we included any trials that were correct and consistent with the old rule in the application phase, and the new rule was marked as starting on the next incorrect feedback.

The following task-based regressors (explanatory variables; EVs) were used in the level-1 model: (1) stimulus presentation (onsets) for rule application trials, (2) onsets for rule learning trials, (3) onsets for trials when participants did not answer, (4) correct feedback, (5) incorrect feedback, and (6) feedback for trials when participants did not answer. These EVs were compared in the following contrasts: (1) rule application versus baseline, (2) rule learning versus baseline, and (3) rule application versus rule learning. Each contrast was tested in both directions (e.g., task > baseline and baseline > task). The baseline (fixation) was implicitly modeled as the intercept, and thus task-based parameter estimates were interpreted as change in activation for task relative to the baseline. Statistical maps describing the results of these contrasts were thresholded to correct for multiple comparisons at the whole-brain level and within an a priori frontal pole mask (ROI), which was defined as the max-probability frontal pole region from the Harvard-Oxford atlas included in FSLView (Desikan, et al., 2006).

To test if activation of the rlPFC may be attributed to difficulty or time-on-task differences, we included a second version of the level-1 analyses, controlling for trial-by-trial reaction times as a separate EV, and testing the same contrasts as described above.

## 3. Results

### 3.1. Behavioral results

To examine how number of rules solved and trials to the rule termination criterion differed between task types, we used a Poisson mixed effects model from the lme4 package (Bates et al., 2015) and a mixed effects Cox regression survival model from the Coxme package (Therneau, 2015) in R (R Core Team, 2014). Consistent with the assumption that the classification task placed more demands on rule evaluation over trials than matching, participants solved more rules during matching (*M* = 15.6, *SD* = 4.14) than during the classification task (*M* = 9.92, *SD* = 3.22), z = 5.58, *p* < .001. Likewise, participants took fewer trials to reach a rule termination criterion in matching (*M* = 6.56, *SD* = 4.67) than in the classification task (*M* = 11.73, *SD* = 7.56), *z* = 12.11, *p* < .001.

To test whether the tasks may have differed in time-on-task, we analyzed the reaction times from correct trials during the application phase with a mixed effects regression model, also from the lme4 package. We found that participants took longer to complete correct matching trials (*M* = 1.78 seconds, *SD* = 0.21) than correct classification trials (*M* = 1.29 seconds, *SD* = 0.26), likely due to the more complex display and larger number of response options, *t*(24) = 9.59, p < .001.

### 3.2. Imaging results

To examine the neuroimaging results, all trials were sorted into either rule learning or rule application phases. In the matching task, all correct trials were considered as rule application trials due to the nature of the task, and all incorrect trials were considered rule learning. In the classification task only the last four or more contiguous correct trials were considered as rule application, and the rest of the trials as rule learning. Trials when participants failed to respond were not included in any phase.

#### 3.2.1. Rule learning and rule application in the matching task

Both the switching and rule evaluation accounts of rlPFC function predict that there should be greater rlPFC activation during the learning phase of the matching task than during the application phase. The switching account predicts greater rlPFC activation during the learning phase because participants are switching between rules during rule learning but not during rule application. The rule evaluation account predicts predicts greater rlPFC activation during the learning phase because participants will know the rule with full certainty and stop evaluating after a single correct answer is found. Consistent with these accounts, we found a cluster in the rlPFC ROI that was activated more for learning than application trials (Figure 6). To verify that this difference was not due to difficulty or time-on-task, we ran a second analysis showing that this cluster remained active when controlling for reaction times (Supplemental Figure 1). No significant activation for the task vs baseline contrasts was observed in rlPFC for the whole-brain or the small-volume corrected ROI analysis.

**Figure 6.**
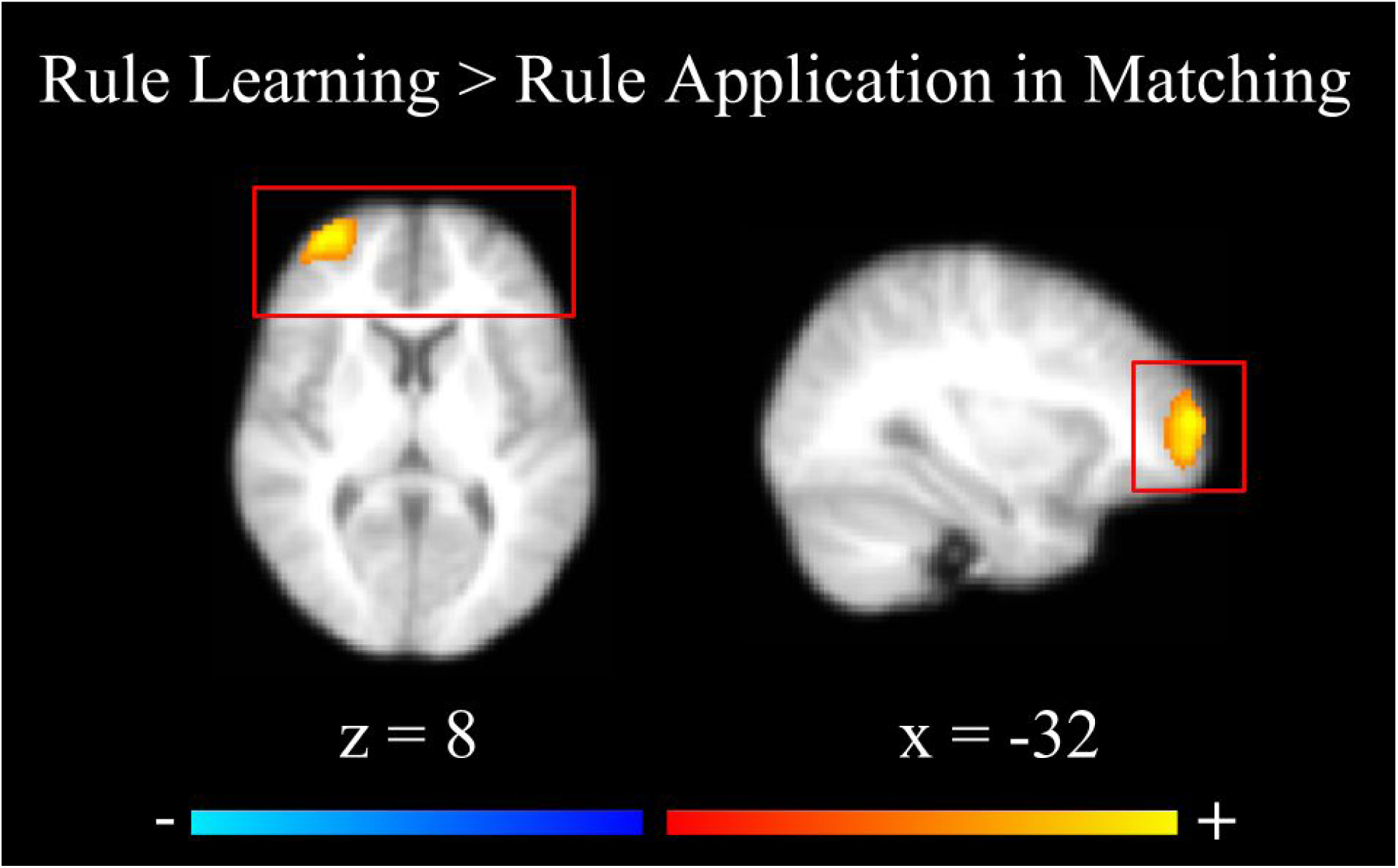
Brain activations in rule learning > rule application contrast in the matching task. The red box is included to convey that the analysis was restricted to a small volume (frontal polar cortex).

Because we did not find any whole brain or ROI-based activation in the rlPFC, to further elucidate the pattern of activation in the rlPFC during the different phases of the matching task, we extracted parameter estimates for each phase from an unbiased ROI based on the classification data. An independent ROI was used to avoid selection bias from the original contrast, which may magnify the size of the difference or bias the direction of activation (Kriegeskorte et al. 2006). This ROI was based on an 8mm sphere drawn around the peak for the rule application versus baseline contrast (MNI coordinates: x = -36, y = 56, z = 4 mm).

As Figure 7 illustrates, the rlPFC is deactivated relative to baseline in the rule application phase for matching, consistent with the possibility that this brain region may cease to contribute to categorization performance once switching or rule evaluation are no longer needed. As with the small volume-corrected ROI analysis, the difference between mean parameter estimates for rule learning and application in the rlPFC was significant, *t*(24) = 3.36, *p* = .002.

**Figure 7.**
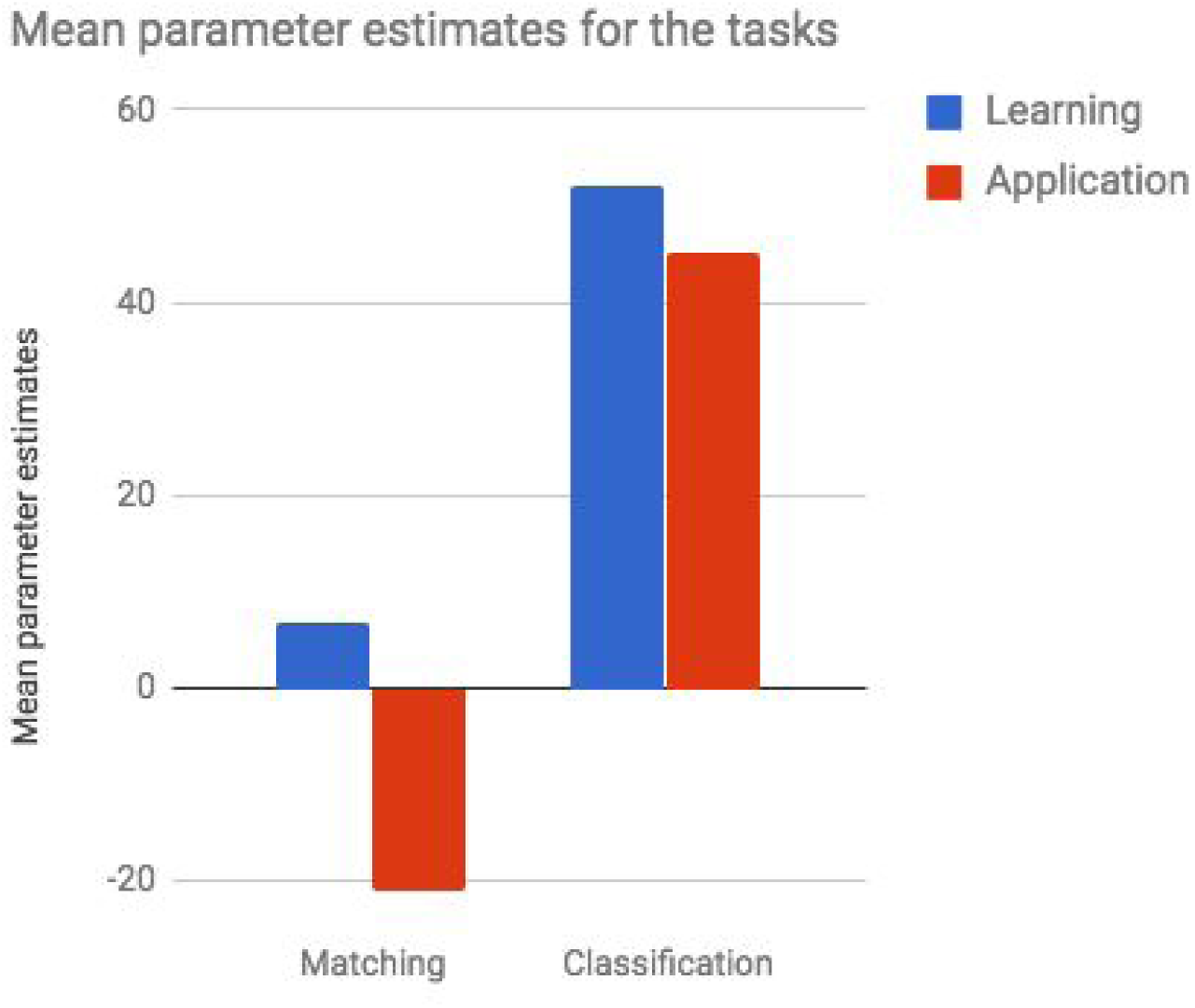
Mean parameter estimates for the matching and classification tasks taken from an rlPFC ROI, where an 8 mm sphere was drawn around the highest peak found in the rule application > baseline contrast for classification task (MNI coordinates: x = -36, y = 56, z = 4 mm).

Beyond the rlPFC, additional areas revealed to be more active during learning than application in the matching task included parietal and lateral occipital regions. These results are consistent with other studies examining activation during learning phases of similar rule-based category learning tasks (Seger & Cincotta, 2006; Liu et al., 2015), and are consistent with previous suggestions that the superior parietal cortex is involved with cognitive control functions during rule-based category learning. Whole-brain results for the opposite contrasts (rule application > rule learning) were also consistent with previous findings from rule-based category learning studies (Seger & Cincotta, 2006; Nomura et al., 2007), with greater activation in the MTL (hippocampus and parahippocampal gyrus) during rule application. Theories of category learning suggest that the MTL is involved in the long-term retention and retrieval of category information during category learning (Ashby & Maddox, 2011; Ashby et al., 2011; Davis et al., 2012a, 2012b). Additional regions more active for rule application included the ventromedial prefrontal cortex, a region we have recently identified as being associated with stronger decision evidence during categorization (Davis et al. 2017), putamen, parietal cortex, occipital cortex, temporal cortex, and insular cortex (all brain maps and other project details are posted on Open Science Framework at https://osf.io/ge8vf/).

#### 3.2.2. Rule learning and rule application in the classification task

In the classification task, we did not find any significant differences in rlPFC activation between learning and application phases in the whole-brain or ROI analysis. Looking further at comparisons between task and baseline, we found that the rlPFC was significantly activated relative to baseline in both rule learning > baseline and rule application > baseline contrasts in both the whole brain and ROI analyses (see Figure 7 for the parameter estimates in rlPFC; see Figure 8 for the whole brain activations in the classification task; also see Supplemental Figure 2 for the whole brain results, controlling for the reaction time). Together, these results suggest that the rlPFC is engaged in a more evaluative role that persists into the application phase in the classification task. This result is inconsistent with switching accounts because participants generally would not be switching rules once they have arrived at a rule that does not result in negative feedback. However, they may continue evaluating confirmatory evidence for their chosen rule because the rule is rarely known with certainty after the first correct trial in the classification phase.

**Figure 8.**
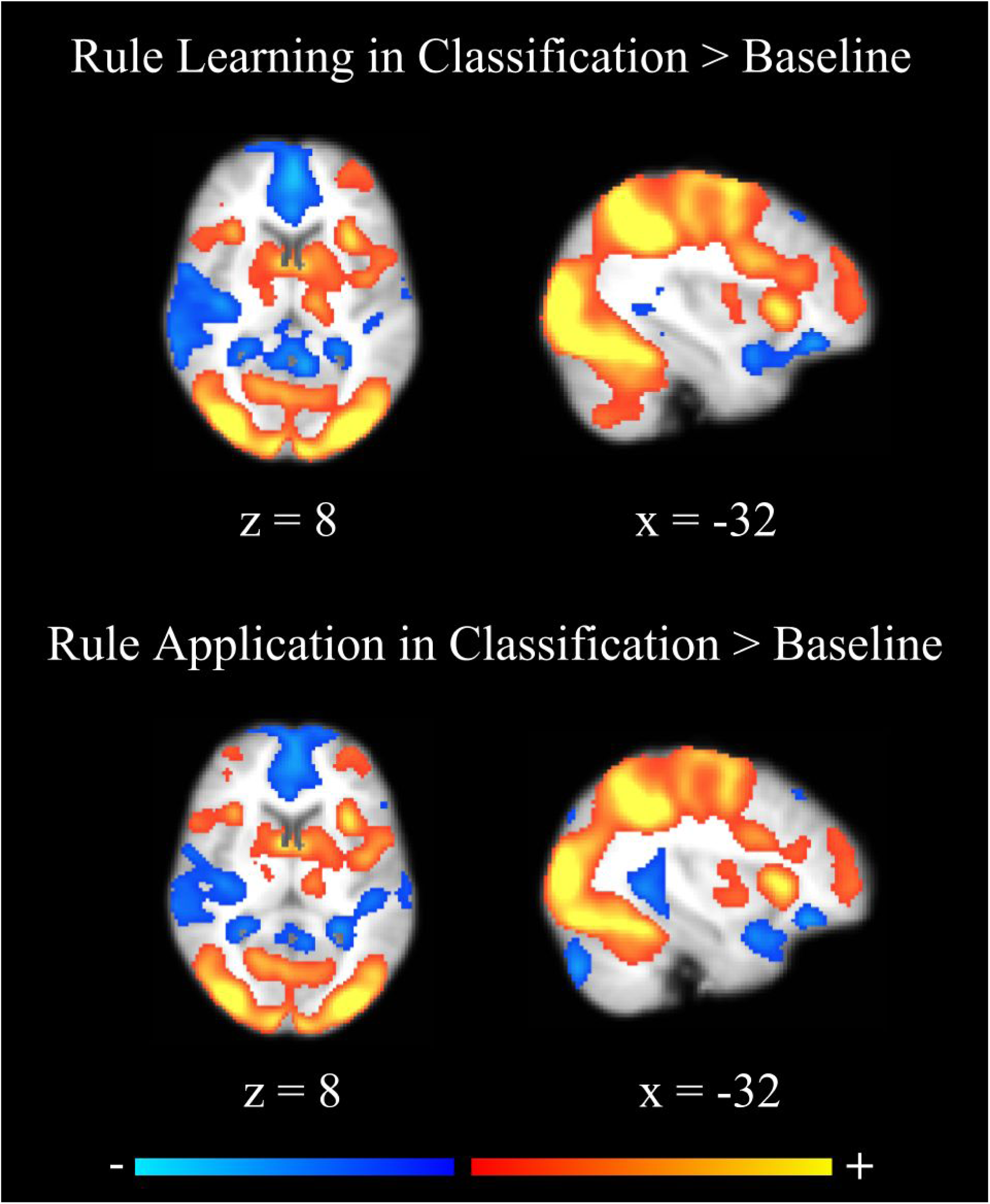
Task vs baseline comparisons for the classification task. Classification > baseline contrasts are in red-yellow and baseline > classification in blue.

Beyond the rlPFC, similar to the matching task, parietal and occipital regions were activated in the rule learning > rule application contrast for the classification task. Additional areas that were more active for rule learning than application included the dorsolateral PFC (see Supplemental Figure 3), which is thought to be involved in maintaining and manipulating information in working memory during rule-based categorization (Monchi et al., 2001; Filoteo et al., 2005; Seger & Cincotta, 2006), and the cerebellum. No regions were significantly more active in rule application compared to rule learning (rule application > rule learning) in the whole brain results.

#### 3.2.3. Comparing rule learning in the matching and classification tasks

The rlPFC was more strongly activated in the classification task than in the matching task during learning (Figure 9) in both whole brain and ROI analyses. These results were not affected by controlling for reaction time, suggesting that they are not due to time-on-task or difficulty differences between the task types (Supplemental Figure 4).

**Figure 9.**
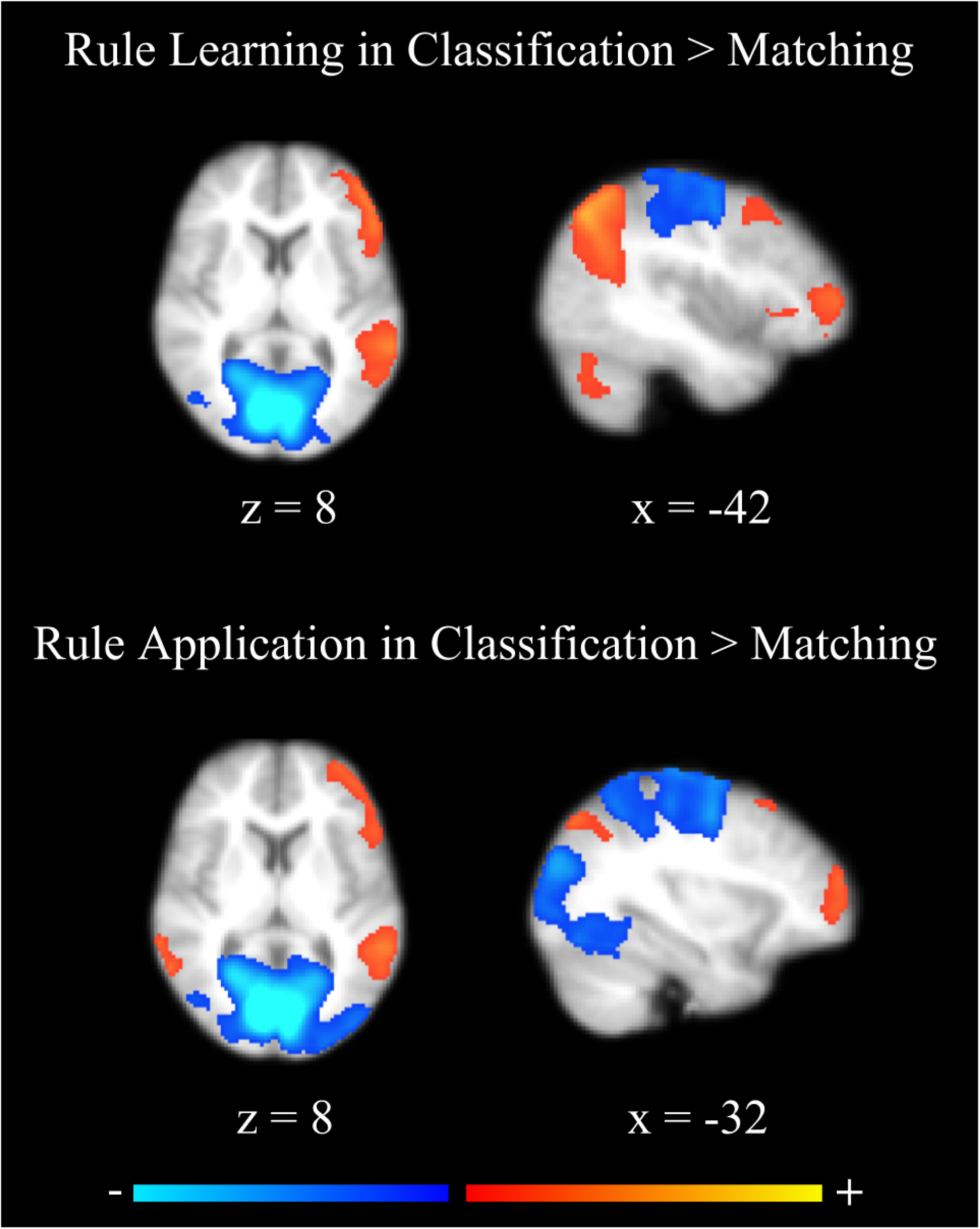
Brain activations for the classification > matching contrasts. Classification > matching contrasts are in red-yellow and matching > classification in blue.

In addition to rlPFC, occipital, inferior parietal, temporal, lateral prefrontal cortices, and cerebellum were more activated for classification compared to matching. Occipital and superior parietal cortices as well as thalamus were more active during the rule learning phase for matching than for classification.

#### 3.2.4. Comparing rule application in the matching and classification tasks

The primary goal of the current study was to compare rule evaluation versus switching accounts of rlPFC function. Because participants quit switching rules during rule application in both matching and classification tasks, but the classification task often involves continued uncertainty and evaluation of evidence for a rule, contrasting rule application during matching and classification allows us to isolate rule evaluation mechanisms. Consistent with the evaluation account, rlPFC was more active in the classification task compared to the matching task. This difference was significant for the whole brain analysis (*p* = .048) and a priori ROI analysis (*p* = .01) in both the main model, and remained significant when controlling for reaction time (Supplemental Figure 4).

In addition to rlPFC, there was greater activation in superior parietal, temporal, and lateral occipital regions in the classification task, and inferior parietal and medial occipital regions in the matching task (see Figure 9 and Table 1 for specific regions covered by the activated clusters). These brain regions overlap with those found in the analyses restricted to the classification and matching phases, and in the analysis examining differences between classification and matching during learning described above.

**Table 1.**
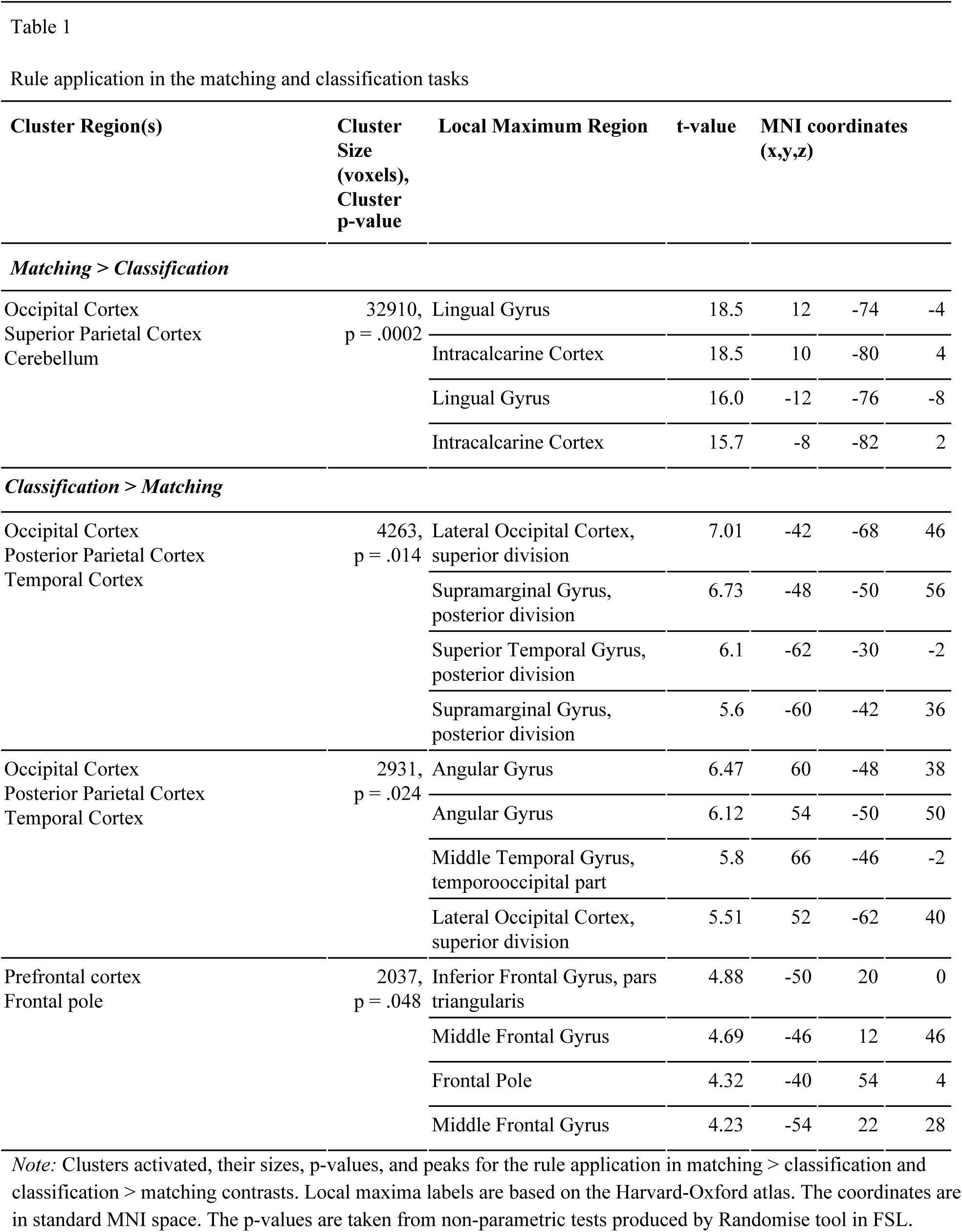
Rule application in the matching and classification tasks

#### 3.2.5. Relationship between rlPFC activation and individual differences in rule application performance

As an additional piece of evidence for the evaluative role of the rlPFC, we also examined how individual differences in rule solving performance related to rlPFC activation in the application versus baseline contrast for classification. We focused specifically on the application phase in classification because it was in this phase that our optimal observer framework suggested continued evaluation demands, but how much continued evaluation demands there were could plausibly be related to how adept individual participants were at solving rules. For this analysis, we extracted parameter estimates from the same rlPFC ROI as described above for the matching task and tested whether they were correlated with individual differences in numbers of rules solved. We found that rlPFC activation was negatively correlated with number of rules solved, *r* = -0.34, *t*(23) = -2.07, *p* = .05 (Figure 10). Given participants with stronger performance should, on average, behave more optimally, and arrive at the rule application phase with less uncertainty and need for continued rule evaluation, this result augments our primary findings and suggests that individual differences in rule evaluation abilities may drive differences in rlPFC activation between subjects. Specifically, the worse participants are at narrowing down rules, the more uncertainty they have when arriving at the rule application phase, and the more they continue to rely on rule evaluation mechanisms in the rlPFC.

**Figure 10.**
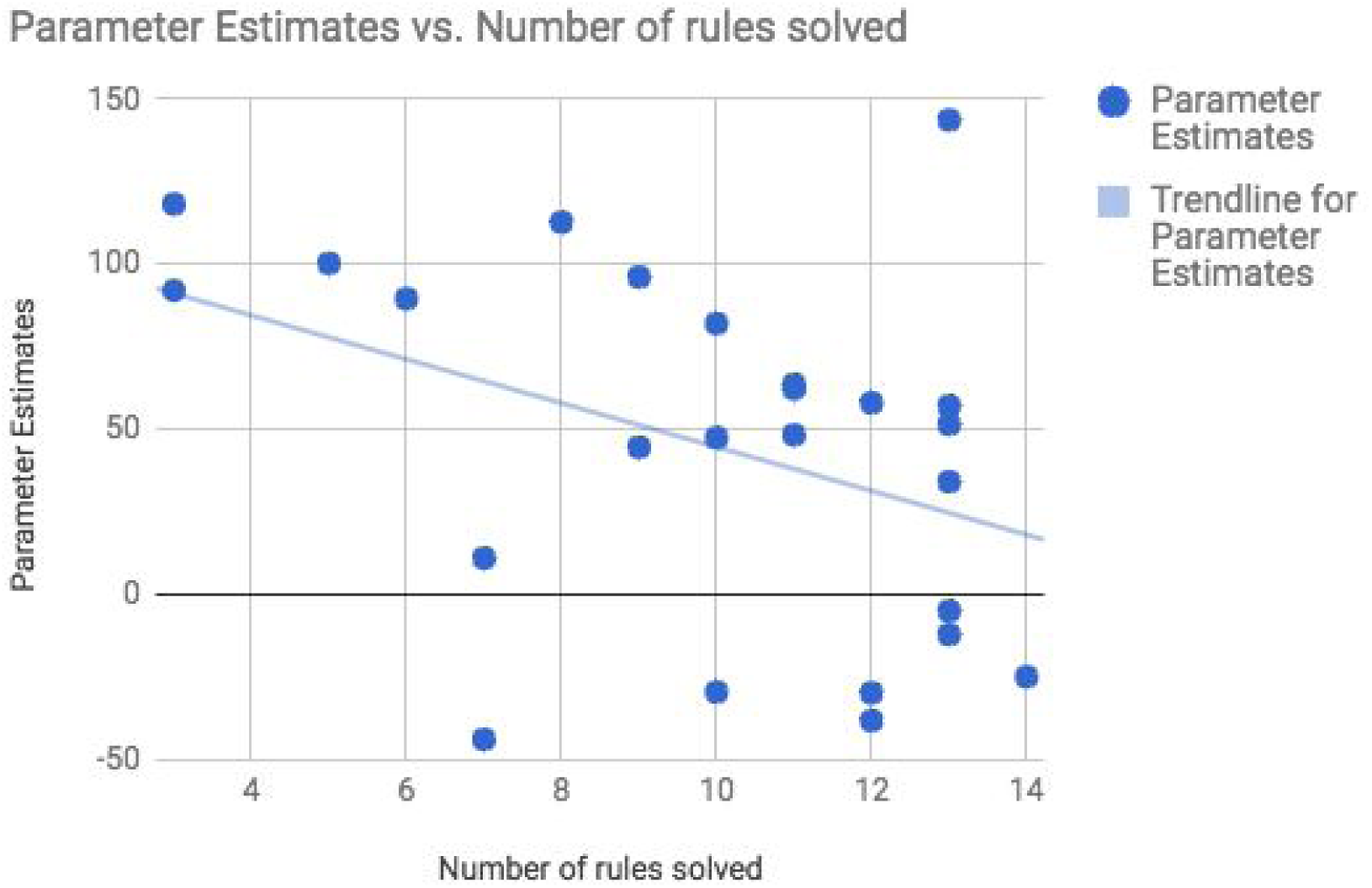
Parameter estimates versus number of rules solved in the classification task. Parameter estimates are taken from an rlPFC ROI, where an 8 mm sphere was drawn around the highest peak found in the rule application > baseline contrast for classification task (MNI coordinates: x = -36, y = 56, z = 4 mm).

## 4. Discussion

The goal of the current study was to compare rule evaluation and switching accounts of rlPFC function in rule-based category learning. The rlPFC has been established as a critical brain region for many higher-order cognitive capacities, yet so far theories of category learning have not fully established a role for the rlPFC. Based on the broader literature, we developed two contrasting accounts suggesting that the rlPFC is involved in rule switching or in the evaluation of evidence for a rule. We tested these contrasting accounts by comparing rlPFC activation during rule-based categorization tasks requiring either matching or classification learning.

Although often treated as the same, these two tasks place different demands on rule evaluation mechanisms. In matching tasks, a rule can be known with certainty after the participant answers a single question correctly and can be eliminated with certainty with every incorrect answer. Therefore participants will tend to evaluate evidence for a rule on the same trials in which they switch between rules. This is not the case in classification tasks, where eliminating a rule can be accomplished in a single trial, but it may be necessary to evaluate evidence over several correct trials before a rule can be established with full certainty. Given this asymmetry in switching and evaluation between matching and classification tasks, we were able to isolate evaluation mechanisms by comparing activation on trials in which participants were applying a rule in matching and classification tasks. Consistent with the hypothesis that the rlPFC is involved in evaluation of evidence for a rule, we found that the rlPFC remained active during rule application in classification learning, but was not active during rule application in matching.

The current results are consistent with other recent studies revealing rlPFC involvement in category learning. Seger and Cincotta (2006) and Liu et al. (2015) found the rlPFC was more active during rule learning compared to rule application in classification-like tasks, and suggested the rlPFC is a part of a “cognitive” cortico-striatal loop involved in rule-based categorization. The current findings are consistent with this hypothesis but go further in suggesting that there may be critical differences between rule-based learning tasks in terms of the demands they place on these rule evaluation mechanisms. This is an important finding as results from matching and classification tasks have often been used interchangeably in the literature on the neural basis of categorization (e.g., Ashby & Maddox, 2005). Our findings suggest that some fundamental contrasts (e.g., rule application vs. learning) can differ markedly between the types of rule-based task employed.

### 4.1. Relationship to multiple learning systems research

By revealing critical differences between types of rule-based tasks, our results fit well within the recent multiple systems literature that focuses on between-task differences in how neural learning systems are recruited (Ell et al., 2006; Nomura et al., 2007; Ashby & Ell, 2001; Ziethamova et al., 2008; Schnyer et al., 2009; for reviews, see Ashby & Maddox, 2005 and Seger & Miller, 2010). However, it is important to note that we are not suggesting that matching and classification involve wholly separate systems, only that they differentially load on these systems, leading to system-level differences in BOLD activation. Future research attempting to build a comprehensive account of the neural basis of categorization will need to take more care in considering commonalities and differences between types of rule-based tasks. Consideration of the demands of particular rule-based tasks may also be critical for neuropsychological assessment where matching tasks, like the Wisconsin Card Sorting Test, are currently popular but may not be as diagnostic of frontal rule evaluation processes as classification learning.

One possible multiple learning systems extension of the current work is to examine differences between the current A/B classification rules (participants choose between category A or B) and rules requiring participants to choose whether stimuli are members or not members of the category (A/not-A). A/not-A rules are mathematically identical to A/B rules (only the labels change), but surprisingly, previous multiple learning systems research has found that these two types of tasks may tap different categorization processes and neural systems (Casale & Ashby, 2008; Zeithamova et al., 2008). Whereas A/B rules seem to recruit brain systems consistent with the episodic memory retrieval, A/non-A may rely more on perceptual memory systems. How A/B vs A/non-A rule types may impact rule evaluation mechanisms in the rlPFC is an open question. One possibility is that A/not-A learning may rely less on explicit rule evaluation mechanisms due to more nonverbal perceptual or procedural strategy use and thus may not recruit rlPFC to the same extent as the A/B tasks we use here.

### 4.2. Relationship to animal learning research

The present results may also inspire future research on rlPFC function in animal models. Recent work on rlPFC function in animal models has demonstrated a critical role for the rlPFC in rapid, one-shot learning of rules, but not in application of well-learned rules (Boschin et al., 2015). Our matching results are consistent with this role of rapid, one-shot rule learning in that the rlPFC was engaged up until participants learned the rule, but then activation dropped to baseline during even early application. However, the classification results suggest that the rlPFC may be engaged even as participants begin to apply the rule, if they need to evaluate additional evidence for the correct rule (e.g., rule out remaining alternatives). Putting these results together suggests that rlPFC can allow one-shot learning, but whether it is engaged for longer-term rule acquisition depends on the demands of the task. Future work with animal models on the rlPFC function would benefit from examining tasks, like our classification task, where evidence for a rule must be accumulated and integrated across trials.

Comparing our study to Boschin et al.’s (2015) results from rule-learning in macaques, one limitation is that we did not have any extended application trials in which participants were applying very well-learned rules for which they had already achieved automaticity. Thus it is not possible, within the current data, to establish whether the rlPFC would remain more engaged for classification learning relative to matching, or if eventually rlPFC involvement would decrease. Given the rapid shift in rlPFC involvement during matching, where it was active during learning but declined in activation as soon as participants began applying the rule, it is likely that rlPFC activation would also decline rather rapidly in classification once the rule was known with certainty. This very brief role for the rlPFC would be consistent with Boschin’s findings from rule learning in macaques as well as other recent studies examining rlPFC involvement in category learning. For example, we recently found, in a relational category learning task (Davis et al., 2017), that rlPFC activation was high early in learning, but in later test phases was re-engaged only when participants needed to generalize the rule to novel relational examples, and not when applying the rule to well-learned examples. In the future, it will be important to test whether the rlPFC exhibits similar trajectories in basic rule-based classification tasks.

### 4.3. Toward a general theory of rlPFC function

Outside of the immediate importance of this work for research on rule-based category learning, the present study adds to a growing literature on rlPFC function in higher-level cognition. Just as it has in category learning, ascribing a single cognitive function to rlPFC has been difficult due to its activation in a wide range of tasks (Duncan & Owen, 2000; Gilbert et al., 2006). For example, the rlPFC has been found to not only be involved in switching (Konishi et al., 1998, 2002; Monchi et al., 2001; Strange et al., 2001) and abstract rule evaluation (Christoff et al., 2001; Kroger et al., 2002; Vendetti & Bunge, 2014; Wendelken et al., 2012; Bunge et al., 2009; Davis et al., 2017; Nee et al., 2014), but also in reinforcement learning (Daw, et al., 2006) and metacognition (Fleming et al., 2012; 2014). Perhaps the best candidate for uniting across these disparate areas of cognition comes from the hierarchical control literature, which suggests that the rlPFC sits atop a rostro-caudal gradient of rule abstraction (Badre, & D'Esposito, 2007; 2009, Badre et al., 2010).

In much of the previous research on hierarchical control theory, participants are given the rules to guide their behavior (e.g., Badre & D'Esposito, 2007)or they learn them in single trials (e.g., in Raven’s-like tasks). In these cases, the minimum abstractness or rule complexity needed to follow or solve the rule is related to which part of the PFC is engaged, with the most abstract rules being processed in rlPFC. Under hierarchical control theory, it is somewhat surprising that rules like those used in our tasks, or metacognitive evaluations of purely perceptual processes (Fleming et al., 2012; 2014), would require the use of the rlPFC. One possibility to reconcile these differences within hierarchical control theory is that the rlPFC is involved whenever participants use rules that involve a structured predicate-argument representation (Ramnani & Owen, 2004; Vendetti & Bunge, 2014), even when these may not be strictly necessary in the task. In the present task, participants may use such representations to guide their initial rule learning and evaluation, but then move back along the control hierarchy to more perceptually based representations after the rules are fully learned and established. Under this hypothesis, the rlPFC may be necessary to learn certain abstract rules, but may be engaged for learning any rule depending on how participants approach it in a given context. Rules based on abstract relations may always depend on rlPFC, whereas rules based on very elementary perceptual features, like the Gabor patches used in Davis et al. (2017) may seldom recruit rlPFC. Between these two extremes may be rules like those in the present task, which can be solved using symbolic representations or perceptual representations (or a combination of both). Whether participants use a symbolic or more perceptual strategy may depend on how easily relevant differences between stimuli are encodable symbolically (e.g., using language; Davis et al., 2009). Although this explanation for integrating the present results with hierarchical control theory is plausible given the broader literature on rlPFC function (Ramnani & Owen, 2004; Vendetti & Bunge, 2014), it remains to be seen whether it can account for rlPFC’s role in other primarily evaluative domains such as in metacognitive judgments of perception (Fleming et al., 2012; 2014).

### 4.4. Limitations and Future Extensions

Relative to some tests of rlPFC function, our study had several limitations in terms of differences in the display and number of options available during the matching and classification tasks. The matching task had four options available (match to the beetle with the same legs, mandibles, antennae, or tail) on every trial, whereas the classification task had only two options (choose category A or B). Likewise, the matching task had more visual clutter on the screen, with four target beetles, whereas the classification task presented only a single beetle at a time.

Although we observed various brain regions associated with visual, motor, and feedback differences between the tasks, the main region of interest, the rlPFC, was found to track predictions from the stimulus evaluation hypothesis both within and between tasks. For example, consistent with the prediction that the rlPFC would not be engaged in the matching task once the rule was known with certainty, rlPFC was engaged during learning but not during rule application in the matching task. Relatedly, consistent with the idea that rule evaluation would persist into the rule application phase in classification, the rlPFC was engaged relative to baseline during both learning and application in the classification task. Because demands were equivalent within task, this pattern where the difference between rule-learning and application was greater in matching than classification cannot be driven solely by button or stimulus display differences. Finally, although theory does not suggest rlPFC should be sensitive to differences in stimulus display or response options, if the rlPFC were sensitive solely to task complexity, the fact that the matching task is more complex in both respects should work against our hypothesis that it would be more engaged during classification.

Although the matching task was the more complex task in terms of response demands and visual complexity, it is possible that other factors like general cognitive difficulty differed between our tasks. Indeed, we expected the classification task to require more rule evaluation demands, and it is possible that activation in some regions reflected increases in general task difficulty caused by these higher demands, as opposed to reflecting the rule evaluation mechanisms themselves. For this reason, we ran additional analyses for all of our main contrasts to control for reaction time, a practice that is used in many fMRI studies to control for potential differences in general cognitive difficulty and/or time-on-task (Brown & Braver, 2005; Grindband et al., 2006; Todd et al., 2013; Davis & Poldrack, 2014; Davis et al., 2014. All of our rlPFC results remained significant, suggesting that general cognitive difficulty is likely not a reason for our observed differences in rlPFC between matching and classification. These results are consistent with findings from the literature on reaction time modeling and perceptual decision making, which do not typically find correlations between measures of difficulty and the rlPFC region that we focused on here (Yarkoni et al., 2009; White et al., 2012; Keuken et al., 2014). For example, in perceptual decision making, measures of decision making difficulty do not typically track activation in the rlPFC (Heekeren et al., 2008; White et al., 2012). Thus taken together, our results are likely to reflect differences in rule evaluation as opposed to general difficulty processing per se. Nonetheless, future studies will want to continue to identify cases where evaluative demands are prima facie fully separated from task difficulty. One domain in which it may be possible to more cleanly separate task difficulty demands from evaluation demands is metacognition, where the rlPFC is often found to negatively correlate with post-decisional confidence (Fleming et al., 2012). Because post decisional confidence ratings are separate from the decisions themselves, they are likely to reflect more pure evaluation signals.

Although differences in task difficulty per se are unlikely to explain our results, there are a number of other factors that may differ between the tasks and should be noted for future research aimed at replicating or extending the current findings. First, from an optimal observer perspective (Figure 1), the classification task may encourage holding more rules in working memory during the application phase of the classification task compared to the matching task.

Indeed, this optimal observer analysis informed our prediction that there would be more remaining uncertainty in the application phase in classification. However, given previous research on how actual participants solve rules in classification tasks that we discuss above (Shepard et al., 1961; Nosofsky, Palmeri, and McKinley, 1993; Wilson & Niv, 2011; Niv et al., 2015), we do not expect participants are explicitly rehearsing more than one rule at a time in either task. Beyond possible working memory differences, however, there remain other differences between the tasks in the total number of trials per rule, the amount of correct or incorrect feedback associated with a rule, and the number of rules completed in each section of the experiment. Although many of these aspects are not expected to affect rlPFC activation given current theory, future research would benefit from attempting to exert greater control over these differences.

Finally, the current study found evidence consistent with the predictions from the evaluation account; The rlPFC’s activation persisted beyond when participants would have been switching rules, suggesting that the rlPFC supports rule evaluation mechanisms. However, it is important to note that these results do not fully establish that the rlPFC is not involved in switching at all. As we discussed above, switching is almost always confounded with evaluation demands. On trials when participants switch rules, they also evaluate the new rule. Thus, while we have provided evidence that the rlPFC is involved in more prolonged rule learning functions, it is not the case that we can fully rule out the possibility that it also participates directly in switching functions beyond its role in evaluation. Fully ruling out switching from uncertainty-related processes like rule evaluation is a major challenge for future research that may be difficult to overcome even using pre-defined, well-learned rules. Switching naturally creates event boundaries in a task, which tend to be associated with higher uncertainty and behavioral variability (e.g., Barcelo et al. 2006; Reynolds et al. 2007). Therefore, it will be important to carefully consider how to create switching situations in the future that are not accompanied by higher evaluative demands and uncertainty.

### 4.5. Conclusion

In conclusion, the present study examines the role of the rlPFC in category learning and differentiates between two accounts of rlPFC function that have been discussed in this literature: rule switching and rule evaluation. To test these accounts, we compared activation during different phases of two types of rule-based tasks, matching and categorization, that differ in their demands on rule evaluation. Consistent with the evaluation hypothesis, the rlPFC was active for tasks requiring more evaluation demands, even when participants were not switching between rules. These results are critical because they help to establish a role for the rlPFC in category learning literature and because they highlight novel differences in the systems engaged for two rule-based tasks that have been used interchangeably in much of the neurobiological literature on category learning thus far. Future research can build on these results by investigating how different types of rule-based representations and task demands impact involvement of rlPFC in rule-based category learning.

## 5. Acknowledgements

We thank Molly Ireland for comments on the manuscript and for proofreading the final version.

